# Intratumoural heterogeneity and immune modulation in lung adenocarcinoma of female smokers and never smokers

**DOI:** 10.1101/2021.05.18.444603

**Authors:** Timo Trefzer, Marc A. Schneider, Katharina Jechow, Lorenz Chua, Thomas Muley, Hauke Winter, Mark Kriegsmann, Michael Meister, Roland Eils, Christian Conrad

**Author notes:** Equal contribution (Timo Trefzer, Marc A. Schneider). Equal contribution as senior and corresponding authors.

## Abstract

Lung cancer is still the leading cause of cancer death worldwide despite declining smoking prevalence in industrialised countries. Although lung cancer is highly associated with smoking status, a significant proportion of lung cancer cases develop in patients who never smoked, with an observable bias towards female never smokers. A better understanding of lung cancer heterogeneity and immune system involvement during tumour evolution and progression in never smokers is therefore highly warranted. We employed single nucleus transcriptomics of surgical lung adenocarcinoma (LADC) and normal lung tissue samples from patients with or without smoking history. Immune cells as well as fibroblasts and endothelial cells respond to tobacco smoke exposure by inducing a highly inflammatory state in normal lung tissue. In the presence of LADC, we identified differentially expressed transcriptional programmes in macrophages and cancer-associated fibroblasts, providing insight into how the niche favours tumour progression. Within tumours, we distinguished eight subpopulations of neoplastic cells in female smokers and never smokers. Through pseudotemporal ordering, we inferred a trajectory towards two differentiated tumour cell states implicated in cancer progression and invasiveness. A proliferating cell population sustaining tumour growth exhibits differential immune modulating signatures in both patient groups. Our results resolve cellular heterogeneity and immune interactions in LADC, with a special emphasis on female never smokers and implications for the design of therapeutic approaches.

## Introduction

The widespread establishment of smoking prevention programs has led to a decrease in the prevalence of smoking, which today accounts for more than 90 % of lung cancer cases [1]. Nonetheless, lung cancer is still the leading cause of cancer deaths worldwide in both men and women, with an age standardised incidence of 18 cases per 100,000 people per year [2-4]. Although many factors contribute to lung cancer susceptibility, a pronounced gender bias among never-smoking patients has been observed. In the year 2000, it was estimated that more than 50 % of lung cancer cases in women worlwide occurred in never smokers, compared to only 15 % in men [5, 6]. Investigating lung cancer with an emphasis on non-smoking patients and especially female never smokers is therefore becoming increasingly important [3, 7-10].

Clinical manifestations of lung cancer are diverse and classification has traditionally been based on histopathological observations. Broadly, lung cancers are divided in two subclasses based on histopathology, non-small cell lung carcinoma (NSCLC) and small cell lung carcinoma (SCLC), with NSCLC accounting for about 80 % of all cases [11, 12]. Within NSCLC, the most frequent subgroup is lung adenocarcinoma (LADC), which is with 60-70 %, also the most prevalent in non-smoking patients [6].

In addition to histological features, recent efforts to stratify patients for improved treatment choices have largely focused on genomic alterations [13], and novel single cell sequencing technologies have dramatically improved our mechanistic understanding of LADC development [14]. This led to the discovery of epithelial transcriptional programs that are deregulated in tumours. A longitudinal study of patients of targeted therapy also identified a transcriptional signature reminiscent of alveolar cells in residual tumour cells during treatment, whereas cells that acquired drug resistance and support tumour progression showed an increase in inflammatory signalling [15]. Single cell studies of LADC have further focused on the tumour microenvironment (TME), providing evidence for changes in cell type composition and transcriptomic profiles in the immune compartment in LADC. Most notably, these included a depletion of immune cells at the tumour site and a bias towards less effective immune cell types, including fewer cytolytic B cells, cytotoxic CD8^+^ T cells, and immunosuppressive macrophages in the tumour vicinity [16-19]. Fibroblast subtypes have been distinguished based on differential expression of e.g. different sets of collagens and endothelial cells showed expression signatures that might contribute to angiogenesis and tissue remodelling [14, 20]. However, the complex interplay of different immune cells and those of the TME remains only partially understood, and therapeutic approaches targeting these cells are still controversially discussed due to intratumour as well as interpatient diversity [21, 22]. Cellular heterogeneity in the immune compartment has profound implications for immune checkpoint therapy, but has proven to be very successful in only a subset of patients. It remains unclear how this heterogeneity, both between patients and within one tumour, might influence therapeutic outcomes for LADC [21-23]. In particular, the role of inflammation as a key immune response contributing to tumour development demands further investigation. Inflammation may be caused by extrinsic factors such as tobacco smoke, as well as air pollution or viral infection, but also by intrinsic processes such as chemokine release, and it promotes cancer initiation and progression by providing an immunosuppressive and tumourigenic environment [24, 25].

Therefore, studying LADC in the context of smoking history with a special focus on female never smoker provides important insights to guide novel therapeutic approaches, which is especially timely with an increasing proportion of LADC cases not attributable to smoking. In this report, we employed single nucleus RNA sequencing to study transcriptional heterogeneity and differentation of LADC tumour cells, as well as the TME, in patients with different smoking habits.

## Results

To investigate the specific characteristics of LADC in never smokers, we retrospectively obtained fresh frozen tissue samples from four patient groups (SupTable. 1 and 2). These included 16 female and 3 male patients between 40 and 60 years of age, to exclude older age as the main confounding factor for any cancer development, and 7 elderly female patients between 75 and 90 years of age. Out of these, 8 of the 16 younger women had a history of smoking while the rest were never smokers. In addition, we included healthy lung tissue from three patients of each group, which we already characterised in a previous publication [26] (Fig. 1A). From all 38 frozen samples, 122,779 intact nuclei were isolated and single nucleus RNA libraries were generated.

**Fig 1.**
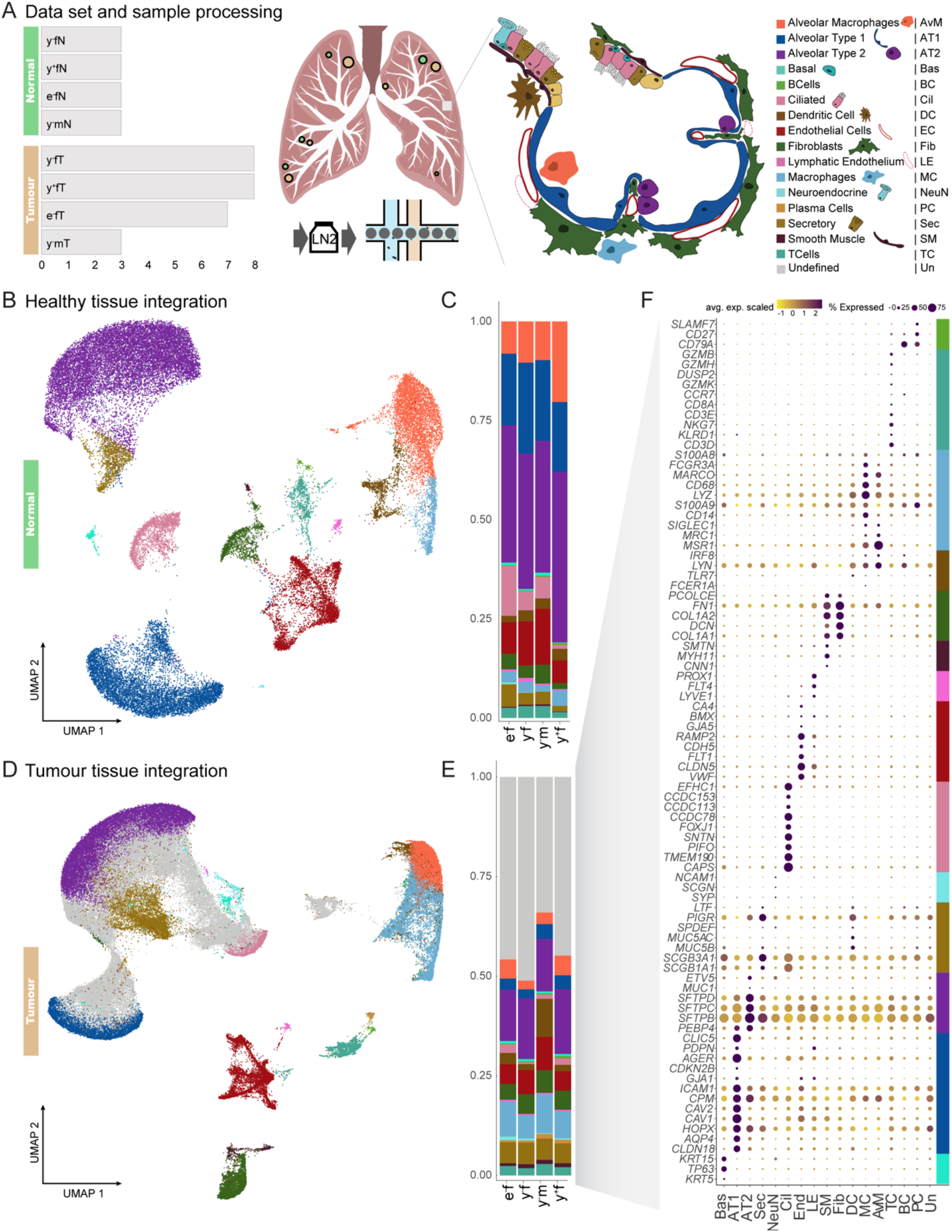
Cell type composition of normal lung and LADC. (**A**) Schematic overview of the samples and cell types of the alveoli. We analysed 38 fresh frozen retrospective surgical samples from 27 patients, including 26 tumour samples (**•**) and 12 matching normal lung samples (**•**) from female (f) and male (m) patients. They comprised two different age cohorts, labelled young (y, 40-60 years) and elderly (e, 75-90 years). Patients are devided in smokers (+) and never smokers (-). Single-nucleus RNA-Seq libraries were prepared from all samples. (**B**) UMAP representation of integrated normal lung transcriptome data. (**C**) Cell type proportions in healthy lung samples. (**D**) UMAP representation of integrated transcriptome data from tumour samples. By comparing gene expression signatures in tumour samples to healthy lung, cell types of the tumour microenvironment (colours as in **B**) and a large fraction of cells with ambiguous expression patterns (grey) were assigned. (**E**) Cell type proportions in tumour samples from the four patient groups. (**F**) Examples of canonical marker gene expression across the cell types identified in tumour samples, with cell type markers indicated by colours as in **A**.

### Smoking leads to an increase in macrophages and decline in endothelial and ciliated cells

We used our previously published atlas of healthy lung single nucleus RNA sequencing (snRNA-Seq) data to serve as a reference for the analysis of single nucleus gene expression in LADC. After integrating data from the 12 healthy lung samples to eliminate technical variation by canonical correlation and mutual nearest neighbour analysis [27], cell types were identified based on differential expression of canonical marker genes (Fig. 1B, SupTable. 3, SupFig. 1E). While cell type composition was broadly comparable between patient groups, smoker lung samples contained fewer ciliated and endothelial cells, increased numbers of macrophages (3.8% in young female smokers compared to 2.7% in never smokers), and more alveolar macrophages (20.4% in smokers compared to 10.4% in never smokers) (Fig. 1C).

Utilizing the established expression patterns of healthy tissue, we inferred cell identities of the tumour microenvironment in all tumour samples (Fig 1D), and found comparable cell type compositions in tumours from all patient groups (Fig. 1E). All cell types identified in healthy tissue were also detected in tumour samples, as verified by their expression of canonical marker genes. In addition, 37,596 cells derived from the tumour samples showed low similarity scores with known lung cell types (SupFig. 1B) and did not specifically express any of the used marker genes (Fig. 1F), suggesting that they represent neoplastic cells.

Our cell type annotation of LADC and healthy lung samples provides a foundation to further characterise the LADC tumour tissue and investigate changes in healthy tissue upon tobacco smoke exposure.

### Inflammatory markers are highly upregulated in female smokers compared to never smokers

Exposure to tobacco smoke causes damage to the lung and elevates cell death, leading to increased infiltration of leukocytes and activation of cellular repair mechanisms [28, 29]. This inflammatory environment is thought to promote lung cancer development and progression. To investigate the effect of smoking in females on gene expression for the diverse lung cell types, we focused on healthy tissue samples of smokers and never smokers between 40 and 60 years of age, thus excluding age and gender as potential confounding factors. Differential expression and gene ontology analysis revealed an enrichment of gene sets relating to inflammation and activation of immune response in smokers (Fig. 2A). Genes involved in these pathways that were upregulated in smokers included *S100A9, SLC11A1* and *NFKB*, which contribute to leukocyte activation and migration, as well as *CCL2, CSF3* and *IL6*, as general mediators of inflammation (Fig. 2B).

**Fig 2.**
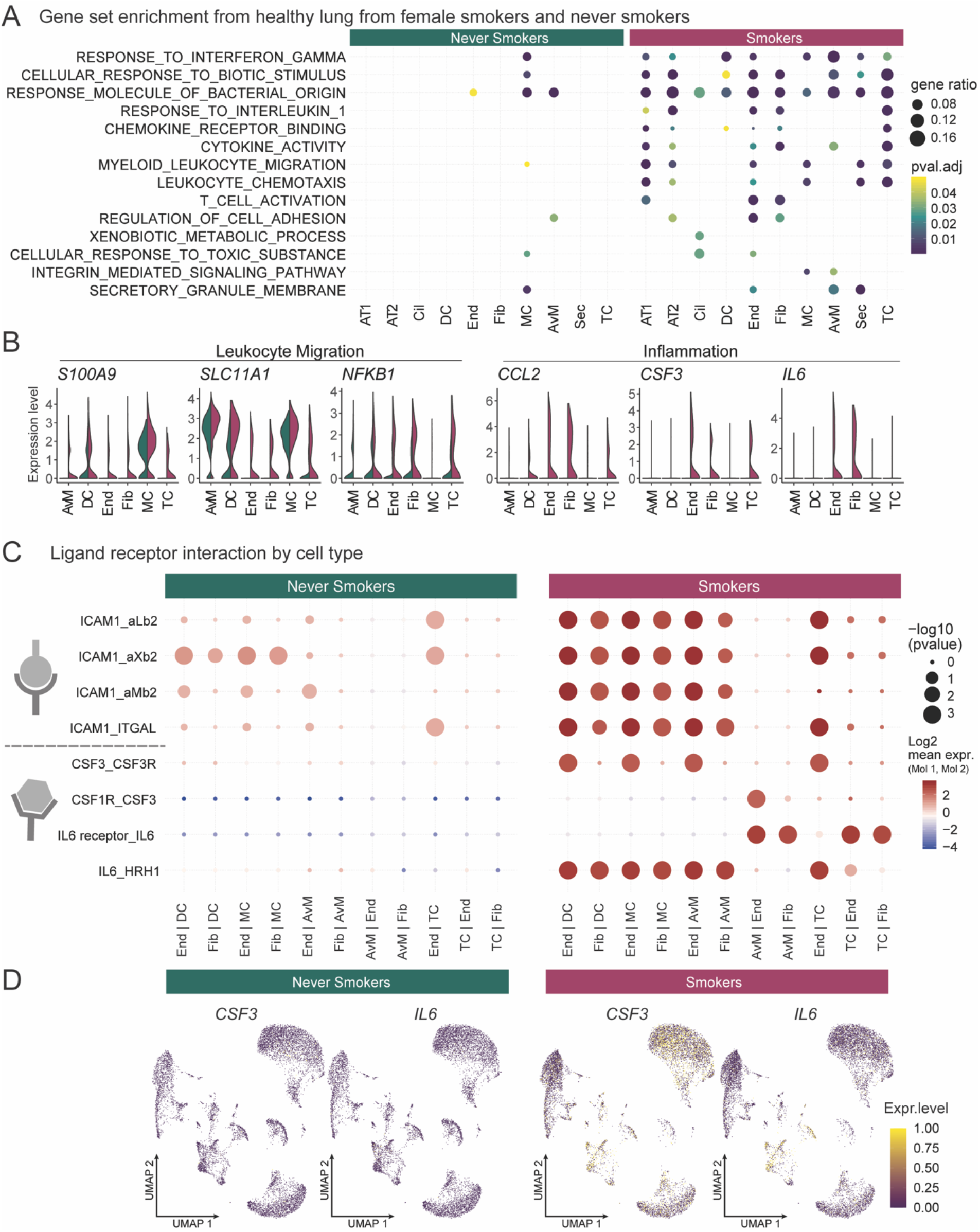
Inflammation and cell interactions in the smoker lung. (**A**) Differential gene expression and gene set enrichment analysis between cells derived from never smokers and smokers by cell type. Celltypes on the x axis as in Fig.1A. (**B**) Expression levels of genes from inflammatory pathways separated by cells from never smokers (green) or smokers (red) samples. (**C**) Putative ligand-receptor interactions inferred from gene expression data for each cell type. (**D**) Expression levels of inflammation mediating ligands *IL6* and *CSF3* in individual cells from never smokers and smokers.

To identify cell type interactions mediating this inflammatory response, we evaluated the expression levels of a curated set of ligand receptor pairs [30] across all cell types, and narrowed down our analyses to cell types with the highest mean expression levels of inflammatory signature genes: immune cell types (dendritic cells, T cells, macrophages and alveolar macrophages) and fibroblasts, endothelial cells and smooth muscle cells. While the overall number of putative ligand receptor interactions is equivalent in smoking and never-smoking patients (SupFig. 2A), interactions of certain drivers of inflammation are increased in smoker lung samples (Fig. 2C). For example, inflammatory cytokines IL6 and CSF3 display an increased expression in endothelial cells and fibroblasts, promoting activation of their corresponding ubiquitous receptors expressed from *IL6R* and *CSF1R* (Fig. 2D and SupFig. 2B). We also identified increased interactions of ICAM1 on endothelial cells, fibroblasts and muscle cells with various integrin complexes on cells of the immune system (Fig. 2D).

Thus, expression changes in fibroblasts and endothelial cells contribute to an inflammatory environment in normal lung tissue in smoking patients, prompting the question whether these differences translate into different tumour phenotypes according to smoking status.

### High intratumoural heterogeneity with distinct cellular subtypes in young female patients

Based on our annotation of tumour sample cells using the healthy lung tissue as reference (Fig. 1D, SupTable. 4), cells that could not be assigned to any endogenous lung cell type were hypothesised to be the transformed cells of the tumour. The deviation from endogenous gene expression signatures in tumour tissue is often caused by mutations or large-scale structural genomic aberrations, such as gains or deletions of chromosomal parts [31, 32]. To corroborate the malignant identity of unassigned cells, we deduced copy number variations (CNV) from transcriptomic data by comparing the average expression levels of genes in close proximity on the genome to a baseline derived from patient-matched normal lung samples (Fig. 3A). Clustering of cells according to their CNV profiles (Fig. 3B) revealed two clusters devoid of copy number variations, which included cells from all patients analysed (cluster 4 and 5), while the other clusters harboured distinct losses or gains and were mostly patient-specific. Clusters containing CNVs were enriched for cells not representative of any healthy lung cell type (Fig. 3B), confirming that these previously unassigned cells are of malignant origin, while the remaining clusters with low CNV prevalence were correctly annotated as cells belonging to the tumour microenvironment.

**Fig 3.**
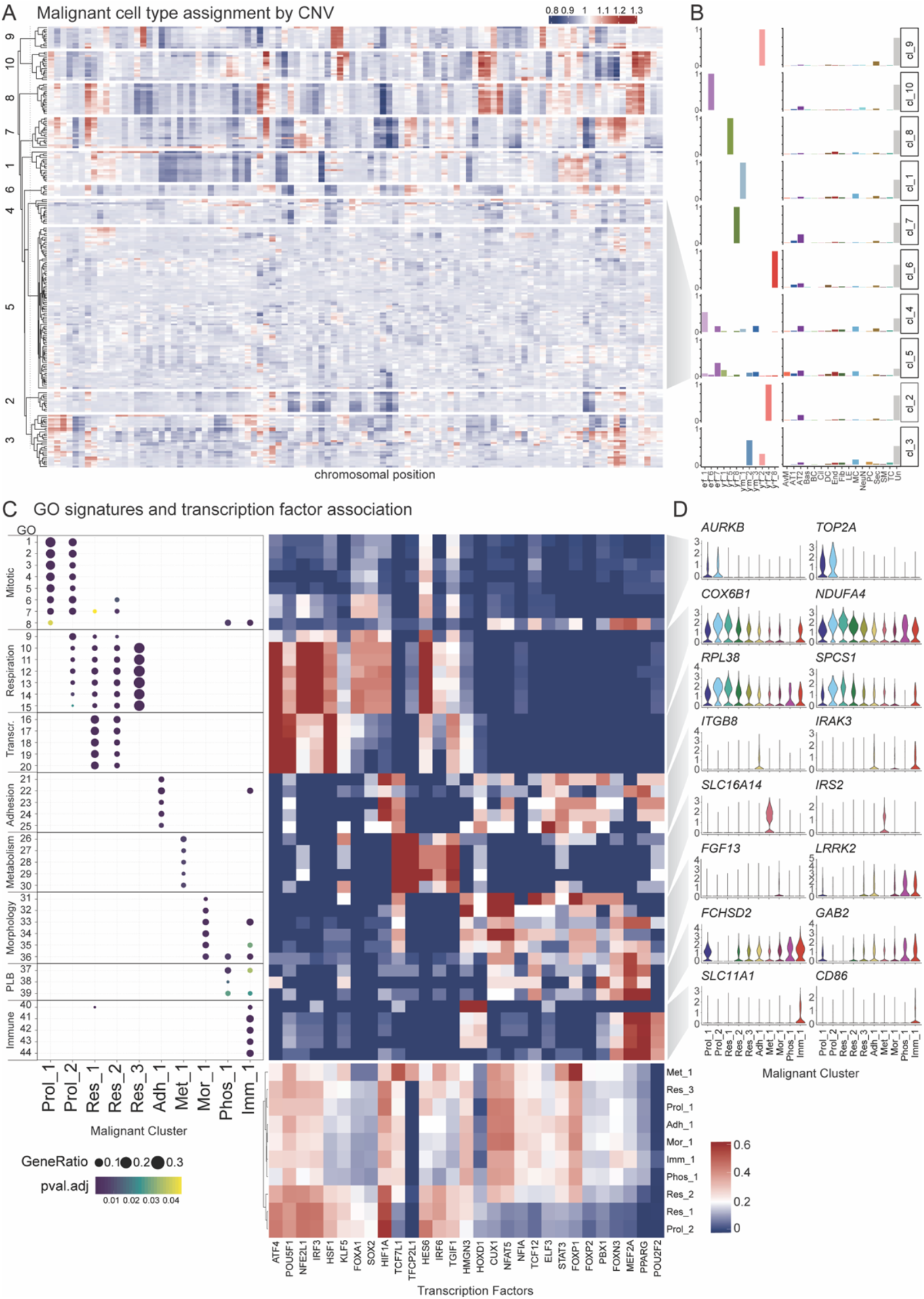
Functional heterogeneity of malignant LADC cells. (**A**) Heatmap of inferred CNV profiles from all cells of tumour samples with matched healthy tissue data. Cell clusters with patient-specific gains (red) or losses (blue) were identified by hierarchical clustering. (**B**) Proportion of cells for each cluster in (**A**) by patient origin (left) with young (y), elderly (e) patients, smokers (+) and never smokers (-), or cell type identity (right) as defined in in Fig.1A. (**C**) Left: Gene set enrichment analysis of differentially expressed genes in malginant cells of females identified 44 cluster-specific GO terms (Supplementary Table 1) that were combined into 8 functional signatures. Right: Transcription factor network analysis of all malignant clusters identified transcription factors contributing to malignant cell cluster identities. Expression level of each transcription factor regulon (bottom) and the fraction of genes associated with a given GO term that are regulated by this transcription factor (top) are shown. (**D**) Expression of representative genes from female patients for each functional signature across malignant cell clusters, named Proliferating (Prol_1/2), Respiration (Res_1/2), Adhesion (Adh_1), Metabolism (Met_1), Morphological (Mor_1), Phospholipid binding (Phos_1) and Immune modulation (Imm_1).

It has been widely demonstrated that solid tumours do not consist of one homogeneous malignant cell population, but represent a heterogenous tissue of diverse cellular states [33]. Clustering of the subset of 37,596 malignant cells from young female LADC patients based on their transcriptome identified ten distinct cell clusters (Fig. 3C, SupFig. 3B) with characteristic gene expression (Fig. 3D). All ten clusters comprised cells from both smokers and never smokers, with some heterogeneity between patients (SupFig. 3C). Enrichment analysis of cluster-specific genes revealed eight expression signatures representative of proliferating cells (labelled ‘Prol’), transcription and cellular respiration (‘Res’), cell adhesion (‘Adh’), metabolism (‘Met’), morphological changes (‘Mor’), phospholipid binding (‘Phos’) and immune related profiles (‘Imm’) (SupTable. 5 and 6). By analysing the expression of transcription factors together with co-expression of their target genes [34], we determined gene regulatory networks contributing to this functional heterogeneity (Fig. 3C). Proliferating cells (Prol) highly express genes linked to networks regulated by ATF4, which is involved in stress responses and amino acid homeostasis [35, 36], and *POU5F1*, also known as *OCT4*, with a critical role in embryonic stem cell self-renewal [37, 38]. Cells enriched for the immune modulating signature (Imm) show additional expression of genes regulated by transcription factors *FOXN3* and *MEF2A*, which are known to be involved in cell cycle checkpoint control and contribute to EMT [39, 40].

Together, these results identify eight functional subpopulations of malignant LADC cells in both smokers and never smokers.

### Trajectory of differentiation and characterisation of malignant cells in context of smoking history

As tumours are evolving and differentiating tissues [41], we applied a graph-based trajectory inference method [42] to malignant cell transcriptomes from young female never smokers and smokers to discern a differentiation trajectory linking the ten functional malignant cell subpopulations identified above. Pseudo-temporal ordering assigned cells to four branches labelled S1-4, with the junction point S0 (Fig. 4A). One branch (S0-S1) consisted of mitotic cells (cluster Prol_1) and immune related signatures (cluster Imm_1) and was therefore selected as the trajectory origin, with pseudotime subsequently increasing through the junction point S0 towards the most distant points on each of the other branches (Fig. 4A). Differential expression and gene set enrichment analysis confirmed cell cycle related gene expression by cells on branch S0-S1, in line with previous findings of cluster signatures (SupTable 7). Branch S0-S4 comprised cells from all identified malignant clusters and was not significantly enriched for specific GO terms; as it was limited to cells at intermediate pseudotimes, some of which were cycling, this branch likely represents undifferentiated tumour cells. Cells on branch S0-S3 largely belonged to cells with morphological changes (cluster Mor_1) and consistently expressed genes involved in cell adhesion, substrate binding and wound healing. Branch S0-S2 mainly harboured respiratory cells (clusters Res_1 and Res_2) with gene expression related to autophagy (Fig. 4B). Together with the respiratory signature of these clusters, this signifies the thight connection between oxidiative phosphorylation and autophagic processes, due to mitochondrial turnover or nutritional need in highly active tissue [43].

**Fig 4.**
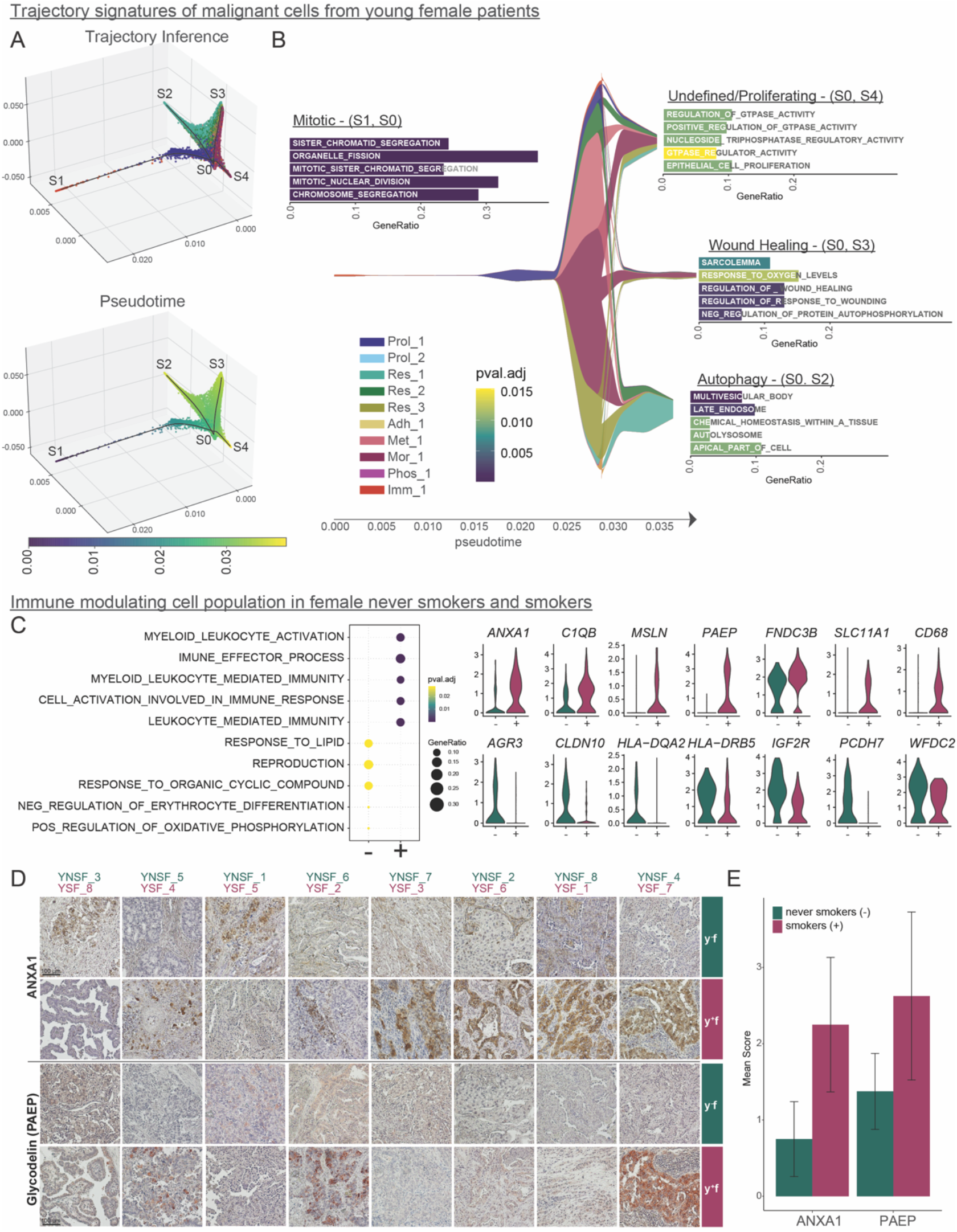
Malignant cell trajectory of female patients. (**A**) Three dimensional projection of cellular expression by Modified Locally Linear Embedding of all identified malignant cells from young female patients to infer a trajectory of differentiation (top) with four branches (S1-4). Cells were ordered by pseudotime (bottom). (**B**) Malignant cluster proportions along pseudotime are depicted. Differential expression and gene set enrichment analysis performed for each branch indicated enrichment of proliferative (S1-S0), undifferentiated (S4-S0), autophagy (S2-S0) or wound healing (S3-S0) signatures. (**C**) Cells from young female patients in cluster Imm_1 were assessed for gene expression differences by smoking habit. Dot plot indicates enriched GO terms in smokers (+) and never smokers (-) Imm_1 cells. Violin plots depict representative genes with significantly different expression levels between never smokers (green) and smokers (red). (D) Immunohistochemical stainings of ANXA1 and glycodelin (*PAEP*) from young female smokers and never smokers. For each patient, one representative staining is shown. (E) Quantification of IHC stainings. Scoring was performed using 5 randomly selected tumor sections combining staining intensities and number of positive cells; displayed are the mean ± s.e.m.

The developmental trajectory thus comprises proliferating and intermediate undifferentiated cells as well as two distinct tumour cell states. Importantly, equivalent trajectories were identified when a separate analysis on malignant cells from young female smokers and never smokers was performed (SupFig. 4), indicating shared functional tumour cell types and a conserved differentiation hierarchy regardless of smoking status.

While LADC from smokers and never smokers in our cohort share the same functional malignant cell types and differentiation trajectory, tobacco smoke exposure might induce more subtle gene expression differences within specific malignant cell types. Comparing gene expression between smokers and never smokers for each malignant cell cluster separately, we observed that the majority of differentially expressed genes were unique to one or two patients, indicating substantial inter-patient transciptional heterogeneity in agreement with the previous analyses (Fig. 3B and SupFig. 1A).

We therefore restricted our attention to genes that were differentially expressed in at least half of the female patients of the same smoking habit, and identified consistent gene expression changes across patients for cluster Imm_1 (Fig. 4C). Here, gene set enrichment analysis uncovered a difference in immune modulating pathway gene expression, with genes including *ANXA1, C1QB* and *PAEP* upregulated in smokers, and genes such as *HLADQA2, HLA-DRB5, WFDC2* upregulated in never smokers. We also observed differential expression of genes involved in migration, EMT and metabolism, with *MSLN* and *FNDC3B* upregulated in smokers and *AGR3, CLDN10, IG2FR* and *PCDH7* upregulated in never smokers (Fig. 4C). To validate gene expression, two exemplary candidate proteins involved in immune modulation pathways were stained in samples from both smokers and never smokers by immunohistochemistry (Fig. 4D). Representative stainings indicate an increased expression of ANXA1 and glycodelin (PAEP) in the majority of female smokers. Quantification of staining intensity revealed a trend for upregulation of both proteins in young female smokers compared to never smokers. Moreover, staining intensity and average gene expression level based on scRNA-seq for each patient correlated for glycodelin, with a trend also observed for ANXA1 (SupFig. 5). This divergence implies differential immune modulating capacity of proliferating tumour cells in female never smokers compared to smokers.

### Tumour microenvironment transcriptome is highly deregulated in LADC

Transformed tumour cells rely to a large extent on interactions with their surroundings, which might either hinder tumour development or work to its benefit [44]. To delineate transcriptomic states within the previously identified cell types of the tumour microenvironment that may contribute to tumour progression, we used non-negative matrix factorisation (NMF) to decompose the gene expression matrix for all non-malignant cell types into the product of two matrices, with the first comprising signatures of co-expressed genes (factors) across all cells and the second capturing the contribution of all genes to these factors (SupTable 8). This approach revealed factors that contribute to cell type identity (Fig. 5A), but also factors that separate cell types into distinct cell states (Fig. 5B).

**Fig 5.**
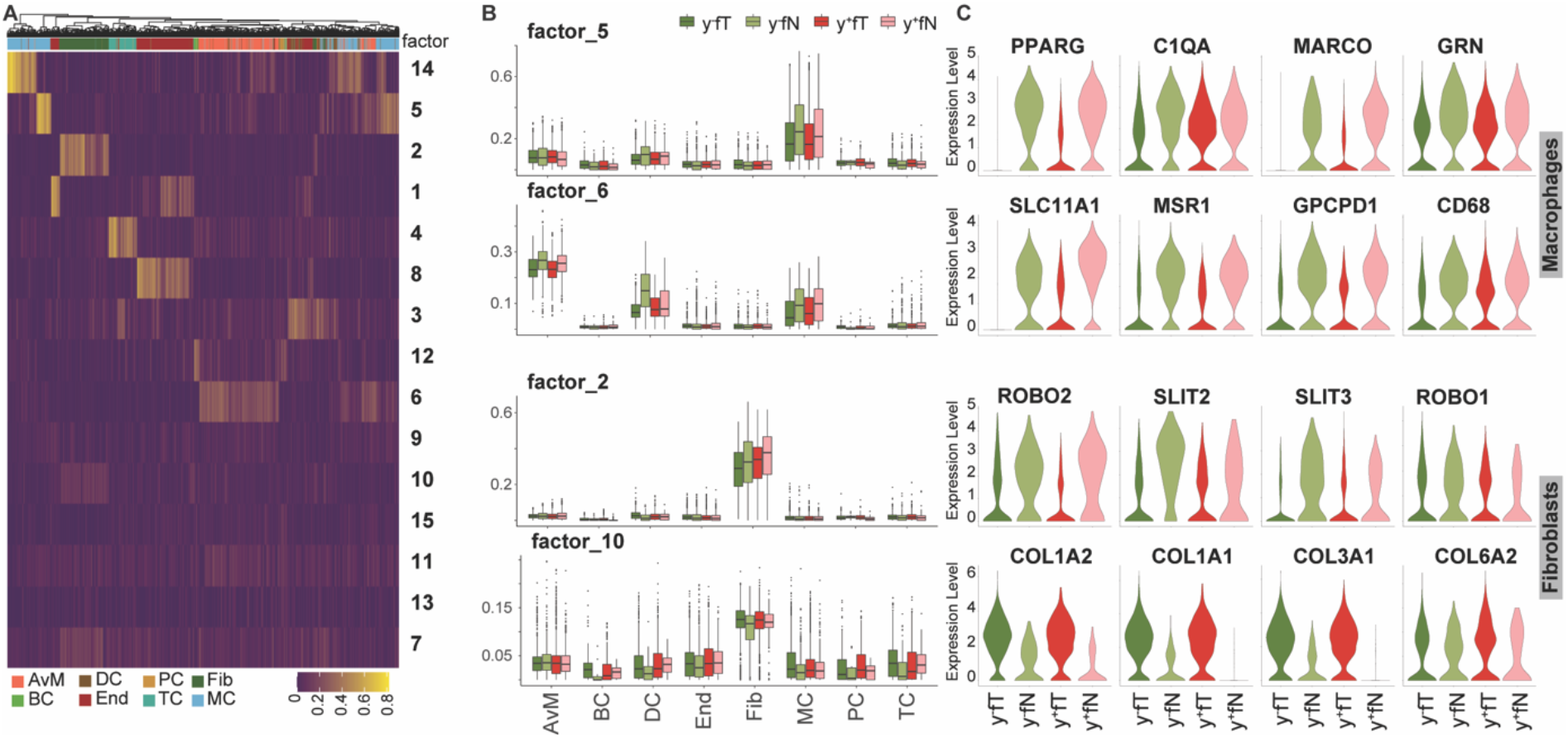
Tumour microenvironment deregulation by smoking. (**A**) Using all non-malignant cells from young female patients in tumour and normal lung samples, gene expression signatures delineating cell types and states were identified by NMF. Color scale indicates factor representation in each cell. (**B**) Contribution of selected factors to observed gene expression in different cell types is depicted separately for tumour and non-tumour tissue samples from never smokers (•/•) and smokers (•/•). (**C**) Expression levels of significant genes represented in the factors shown in (B), across tumour and non-tumour tissue samples from never smokers (•/•) and smokers (•/•).

Two of these factors (factor 5 and 6) represent two cell states within the macrophage population with decreased expression in tumour tissue compared to healthy lung (Fig. 5C), and contain genes involved in immune cell activation and inflammation (e.g. *PPARG, C1QA, MARCO, GRN* and *SLC11A1, MSR1, GPCPD1, CD68*). Specifically, factor 5 contains genes with a role in macrophage activation such as the ones encoding for PLXDC1, a receptor of ligand PEDF that enhances tumouricidal activity of macrophages [45, 46], and SLC11A1, a divalent transition metal transporter whose activity is associated with pro-inflammatory processes [47]. Downregulation of this signature indicates a reduced activation of macrophages in the presence of LADC. Factor 6 delineates another subpopulation of macrophages with lower expression of inflammatory genes in tumour tissue, including *PPARG* and *MARCO*. The latter has been suggested as a possible treatment target in NSCLC [48], since antibody targeting of MARCO expressing macrophages reduced tumour growth in a recent study [49]. Consistent with our observation that only a subset of macrophages downregulate *MARCO*, the same study found MARCO expression in only a subset of tumor-associated macrophages. Anti-MARCO antibody treatment was therefore most effective in combination with other immune checkpoint markers [49].

Along with macrophages, fibroblasts can also support or hinder tumour development. We identified a population of fibroblasts with decreased expression of genes in the *SLIT*/*ROBO* pathway in tumour samples (factor 2). The *SLIT*/*ROBO* pathway has often been found to be differentially regulated in cancer, where its complex involvement in tumour progression may include beneficial as well as detrimental effects on tumour growth [50]. The observed decreased expression of *SLIT2* may facilitate tumour survival and cell progression [50], while *SLIT3* downregulation might enhance epithelial to mesenchymal transition (EMT) [51]. Another population of fibroblasts showed increased expression of type I and type III collagens in neoplastic tissue (factor 10). As part of the tumour microenvironment, different extracellular matrix components provided by fibroblasts have been found to affect tumour behaviour [52]. Increased expression of type I and type III collagens, as observed here, is thought to promote invasion and metastasis in lung cancer [53-55].

Our results thus resolve different macrophage and fibroblast subpopulations in LADC, with distinct gene expression signatures contributing to a tumourigenic environment in both smokers and never smokers.

## Discussion

In this study, we employed single nucleus RNA sequencing of fresh frozen surgical tumour samples of LADC, together with patient-matched healthy lung samples, to investigate cellular heterogeneity, cellular interactions and the TME in patients with or without a smoking history. In worldwide never smoker lung cancer cases there exists an observable bias towards women [5]. We therefore focused this study on female smokers and never smokers.

Tumour cell heterogeneity is increasingly recognised to play a crucial role in tumour progression, with implications for tumour evolution and efficacy of treatments [15, 56]. Within the LADC malignant cell compartment, we resolved distinct cell populations with eight gene expression signatures representing cell proliferation, cellular respiration, transcription, cell adhesion, metabolism, morphological changes, phospholipid binding and immune modulation pathways. Linking these signatures to transcription factor expression also indicated gene regulatory networks contributing to the observed functional heterogeneity. By ordering malignant cells along a pseudotemporal trajectory, we inferred a progression of cell states from proliferating cells, via an undifferentiated state that comprised cells from the majority of malignant cell types, towards two distinct endpoints representing signatures of either autophagy or wound healing processes. Wound healing mechanisms have long been suggested to be involved in cancer progression, invasion and metastasis by creating a niche that fosters proliferation and tissue remodelling [57-61]. The detection of autophagy signatures may reflect the struggle or development of cells in this trajectory as autophagy has an ambiguous role in cancer progression with implications for mitochondrial turnover in metabolically highly active cells and indicator for nutrient defficency in poorly vascularised tissue [43, 62, 63].

Furthermore, we resolved specific gene expression signatures that were downregulated in macrophages in the LADC tumour microenvironment compared to healthy lung, and two populations of cancer associated fibroblasts with differential expression of genes involved in cell migration, angiogenesis and metastasis. These expression signatures could aid in the design of therapeutic approaches that target the TME.

Interestingly, no distinct cell populations unique to female smokers and never smokers were detected in either normal or malignant tissue samples, and tumour developmental hierarchies were equivalent across patient groups (Fig. 4, SupFig. 3 and SupFig. 4). However, taking into account the profound inter-patient heterogeneity (Fig. 3A/B and SupFig. 1A), we resolved distinct transcriptional properties of the cluster of malignant immune modulating cells (Imm_1) according to smoking history, with increased expression of immune-related genes such as *ANXA1, C1QB, SLC11A1, CD68, PAEP* in smoker cells and *HLA-DQA2, HLA-DRB5, WFDC2* in female never smokers. In the same cell cluster, we also identified genes involved in migration and development that were specifically expressed in smokers (*MSLN, FNDC3B*) or never smokers (*AGR3, CLDN10, IGF2R, PCDH7*). We validated the increased expression of ANXA1 and glycodelin (*PAEP*) by immunohistochemistry. Annexin A1 (ANXA1) inhibits cytosolic phospholipase A2 (PLA2) and is therefore considered as an anti-inflammatory agent [64]. Moreover, ANXA1 expression has been shown to be elevated in patients with COPD [65], but downregulated in female never smokers with NSCLC [66]. Glycodelin has been well characterized concerning its immunosupressive function at the fetomaternal interface [67, 68], but has also received increasing attention as an immunomodulatory marker for cancers including melanoma [69] and NSCLC [70] over the last decade. The subset of proliferating cancer cells might therefore differentially modulate the immune microenvironment according to patient background and smoking status, with potential significance for immunotherapies.

In addition to gene expression differences within the malignant cell compartment, our analysis of normal lung tissue samples identified an increase in inflammation and immune activation induced by tobacco smoke exposure, with inflammatory signalling molecules such as CSF3, ICAM1 and IL6 mediating communication between immune cells as well as fibroblasts and endothelial cells (Fig. 2). This was consistent with a decreased proportion of ciliated and endothelial cells and higher proportion of macrophages in smoking patients (Fig. 1C), indicating that inflammation may lead to invasion of macrophages and tissue damage. Similar consequences have been proposed based on histology, lavage, elevated inflammatory molecules in peripheral blood or bulk transcriptome samples [28, 29, 71-73]. While more recent single cell transcriptomic studies investigated the effects of tobacco smoke in systemic immune cells and upper airway epithelial cells [74-76], cell types in the alveolar region and their interplay had not been addressed at this resolution. Our results might thus aid in the identification of therapeutic agents that could counteract the known tumorigenic effects of inflammation, a challenge that remains unsolved [77, 78].

A significant obstacle in the analysis of tumour tissues within this study was the substantial inter-patient heterogeneity, as previously observed. Due to differences in genetic background, epigenetic modifications, patient history and comorbidities, this can only partly be overcome by larger sample sizes and molecular patient stratification. Computational methods that exclude patient specific features without losing biologically relevant signals will be necessary to further refine analyses of malignant cell populations across patients at the single cell level. In addition, our results based on single cell transcriptomics could be tested in larger patient collectives using bulk omics approaches. As smoking prevalence decreases, future studies should also address other environmental and intrinsic factors contributing to inflammation. These include inflammatory diseases such as chronic obstructive pulmonary disorder (COPD), which increases the risk of lung cancer independent of age, sex and smoking status [79, 80].

In conclusion, we here identify key cell types and pathways contributing to the highly inflammatory environment in smoker lungs. Analysing the especially susceptible group of female never smokers, we provide a refined description of cellular heterogeneity within LADC tumours and their microenvironment, and define transcriptional signatures for distinct transformed cell states. While the cell type composition and differentiation hierarchy of LADC were equivalent in female smokers and never smokers, we identified a subset of cells with differential immune modulating activity dependent on smoking status. These findings will aid in the selection and development of treatments that take into account the complex interplay of disease aetiology, intratumoural heterogeneity and interactions with the tumour microenvironment.

## Supplementary Materials

**SupFig 1.**
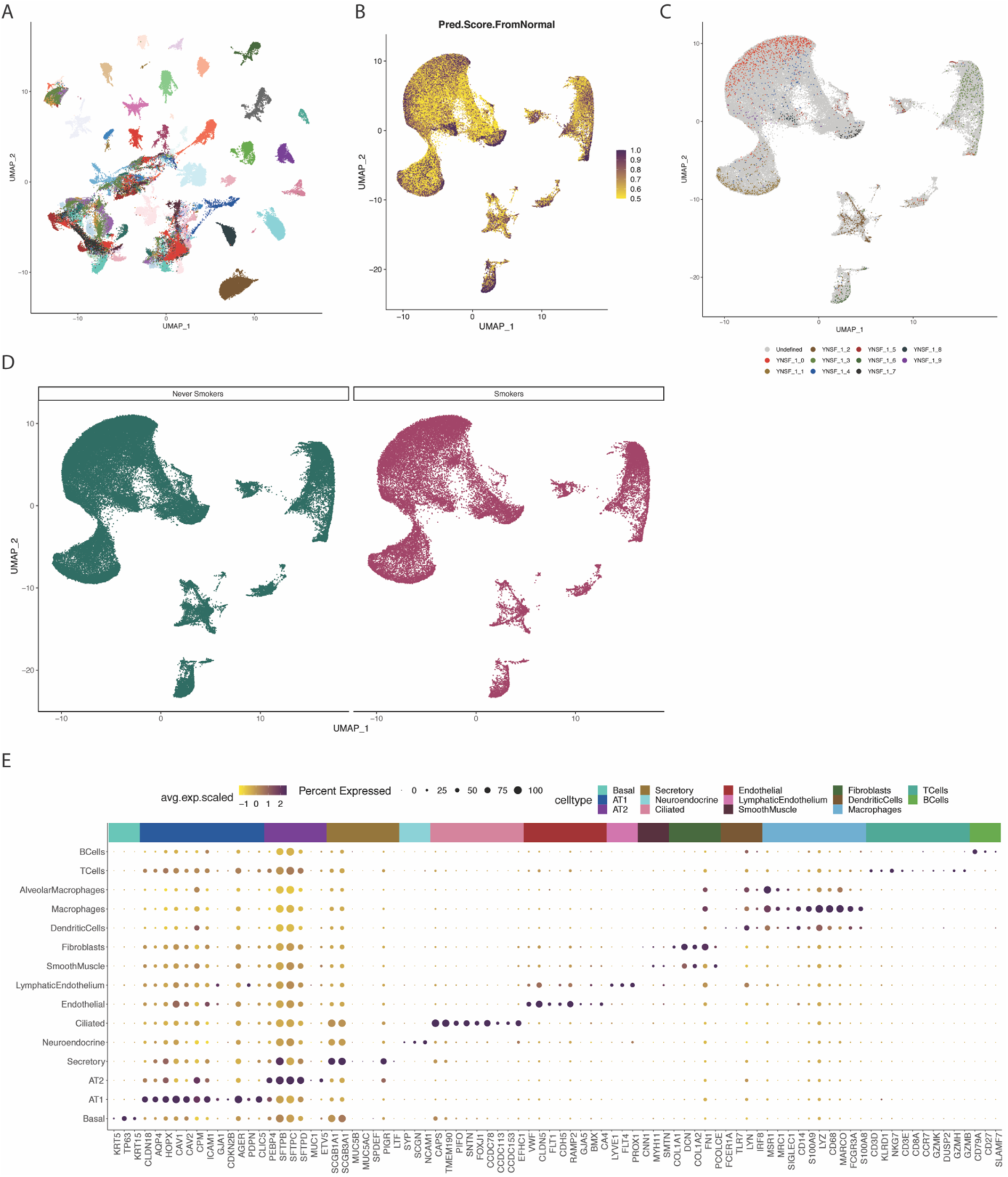
Data integration. (**A**) UMAP representation of single-nucleus transcriptomes from all samples prior to integration by CCA. (**B**) Cell types were assigned for cells originating from tumour tissue by comparing gene expression signatures to the healthy lung reference data. Depicted are confidence scores for the assignment. (**C**) Clusters identified through separate analysis of transcriptome data from a single patient are projected on the integrated UMAP of all samples. Integration did not disrupt cluster identity. (**D**) Cells are separated by the smoking habit of patients, showing no cluster unique to either patient group. (**E**) Expression of representative marker genes in healthy lung reference samples across all annotated cell types.

**SupFig 2.**
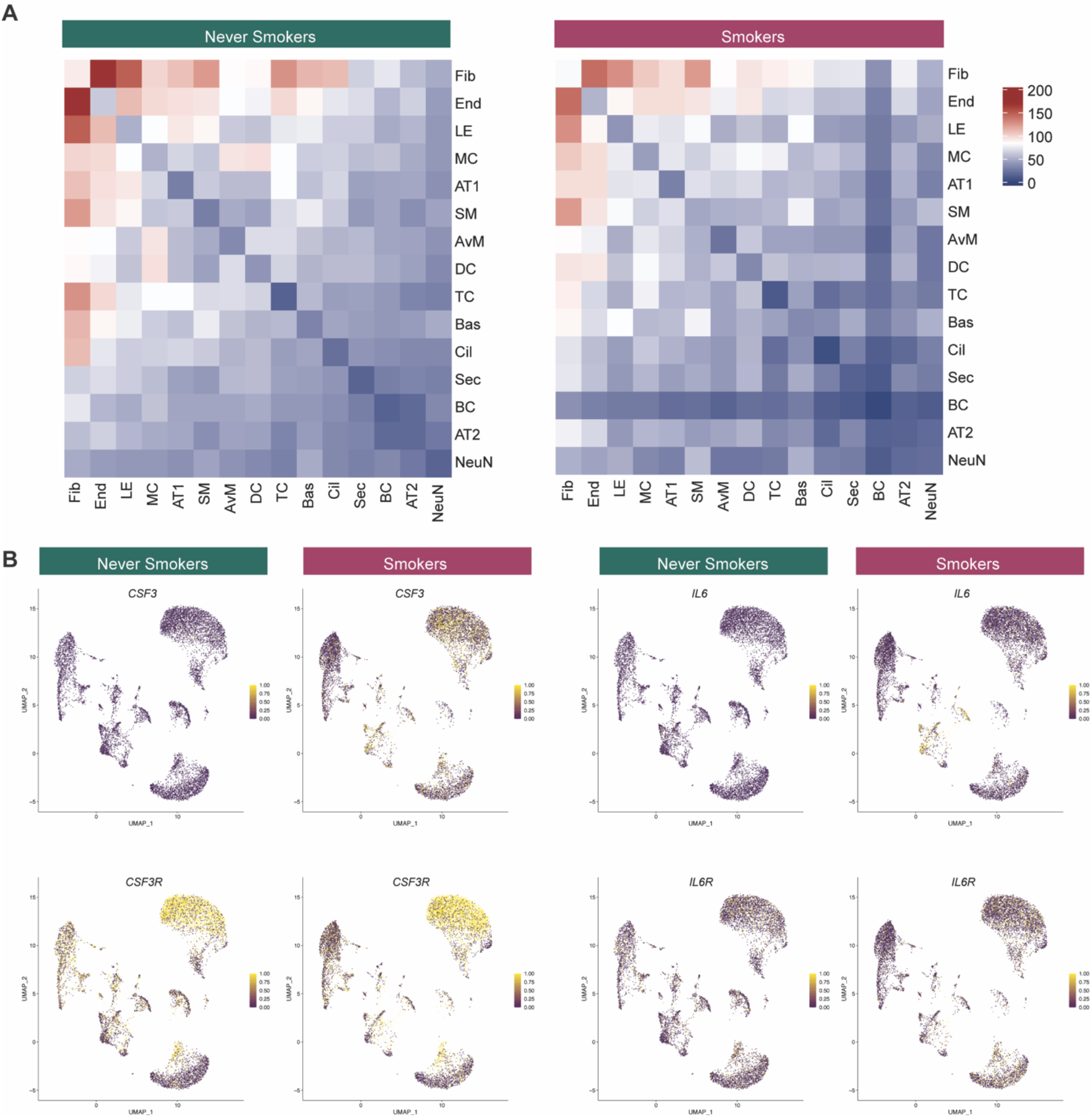
Ligand receptor interactions. (**A**) Overall number of inferred cell to cell interactions in the different annotated cell types, showing no significant difference by smoking habit. (**B**) Ligand and receptor gene expression of selected chemokines across all non-malignant cells. UMAP representation is the same as in Fig. 1B.

**SupFig 3.**
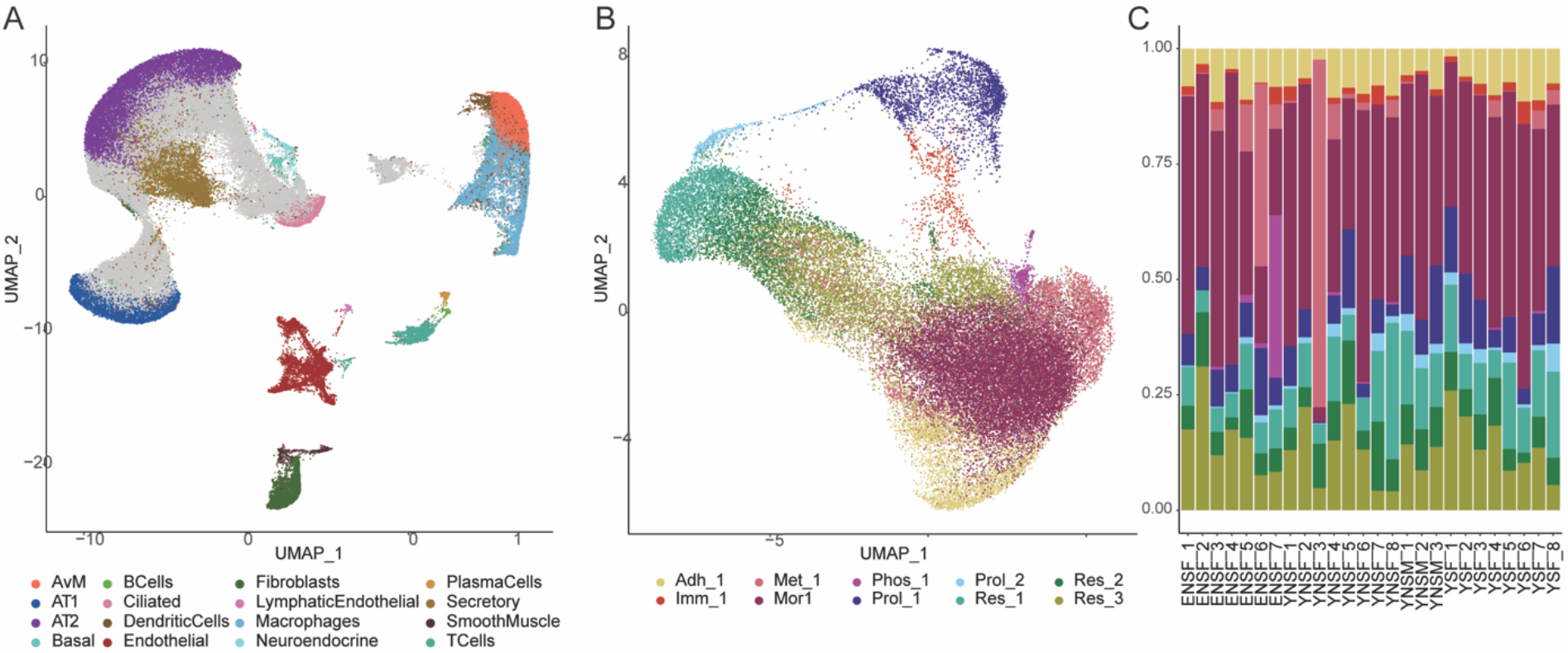
Malignant cell heterogeneity. (**A**) Integrated UMAP representation of all cells including tumour and healthy tissue samples. (**B**) UMAP representation of cell clusters within the malignant cell compartment, as identified by CNV analysis. (**C**) Proportions of malignant cell clusters by patient. Elderly never smokers female (ENSF), young never smokers feamle (YNSF), young smokers female (YSF) and young never smokers male (YNSM).

**SupFig 4.**
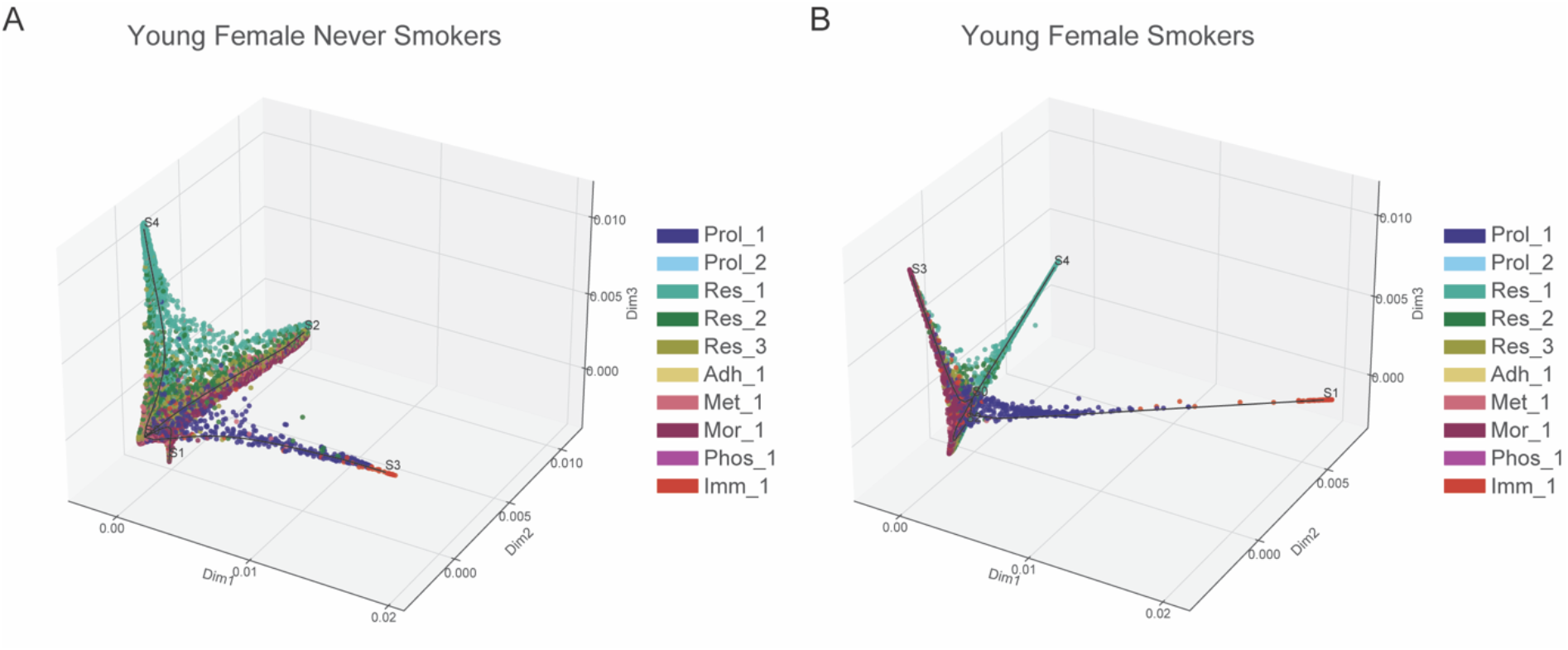
Malignant cell trajectory. Trajectory inference of malignant cells from only (**A**) young female never smokers or (**B**) young female smokers.

**SupFig 5.**
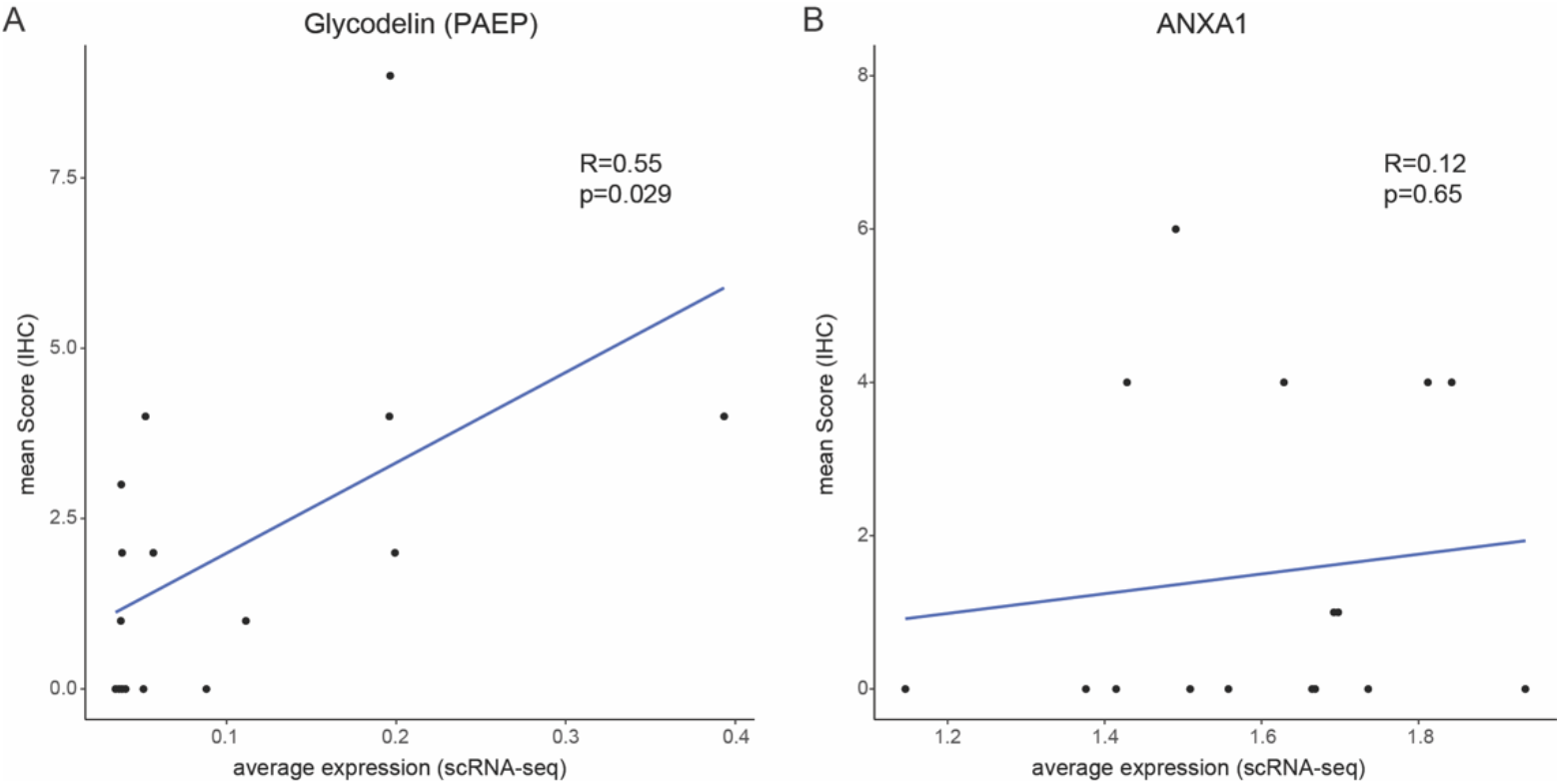
Correlation of IHC and scRNA-seq expression of PAEP and ANXA1. Correlation of protein expression determined by quantitative scoring of immunohistochemistry staining and average gene expression for (**A**) Glycodelin (PAEP) and (**B**) ANXA1.

**Supplementary Table 1 –.** Patient cohort overview.

| **Cohort description** |  |  |
| --- | --- | --- |
| **Parameter** | **n** | **(%)** |
| Median Age | 52 (40–88) |  |
| **Total** | **26** | **100** |
| Male | 3 | 12 |
| Female | 23 | 88 |
| **Histology** |  |  |
| Adeno | 26 | 100 |
| **Therapy** |  |  |
| OP | 15 | 58 |
| OP/RT | 1 | 4 |
| OP/ChT | 9 | 34 |
| OP/RT/ChT | 1 | 4 |
| **Smoking status** |  |  |
| Never Smokers | 18 | 69 |
| Smokers | 8 | 31 |
| **Pathological Stage**<br>**(7<sup>th</sup> TNM edition)** |  |  |
| IA | 2 | 8 |
| IB | 9 | 35 |
| IIA | 4 | 15 |
| IIB | 2 | 8 |
| IIIA | 7 | 27 |
| IIIB | 2 | 8 |
| **ECOG** |  |  |
| 0 | 26 | 100 |

**Supplementary Table 2 –.**
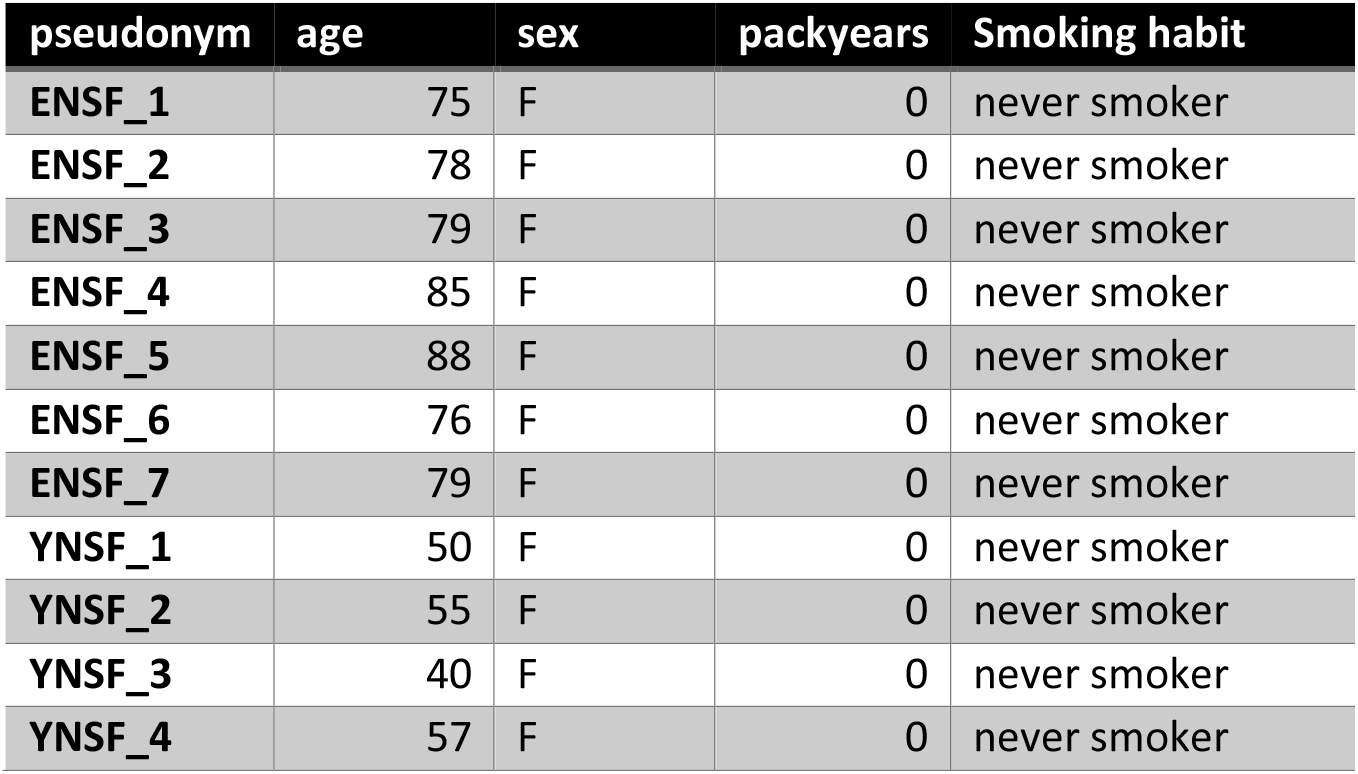
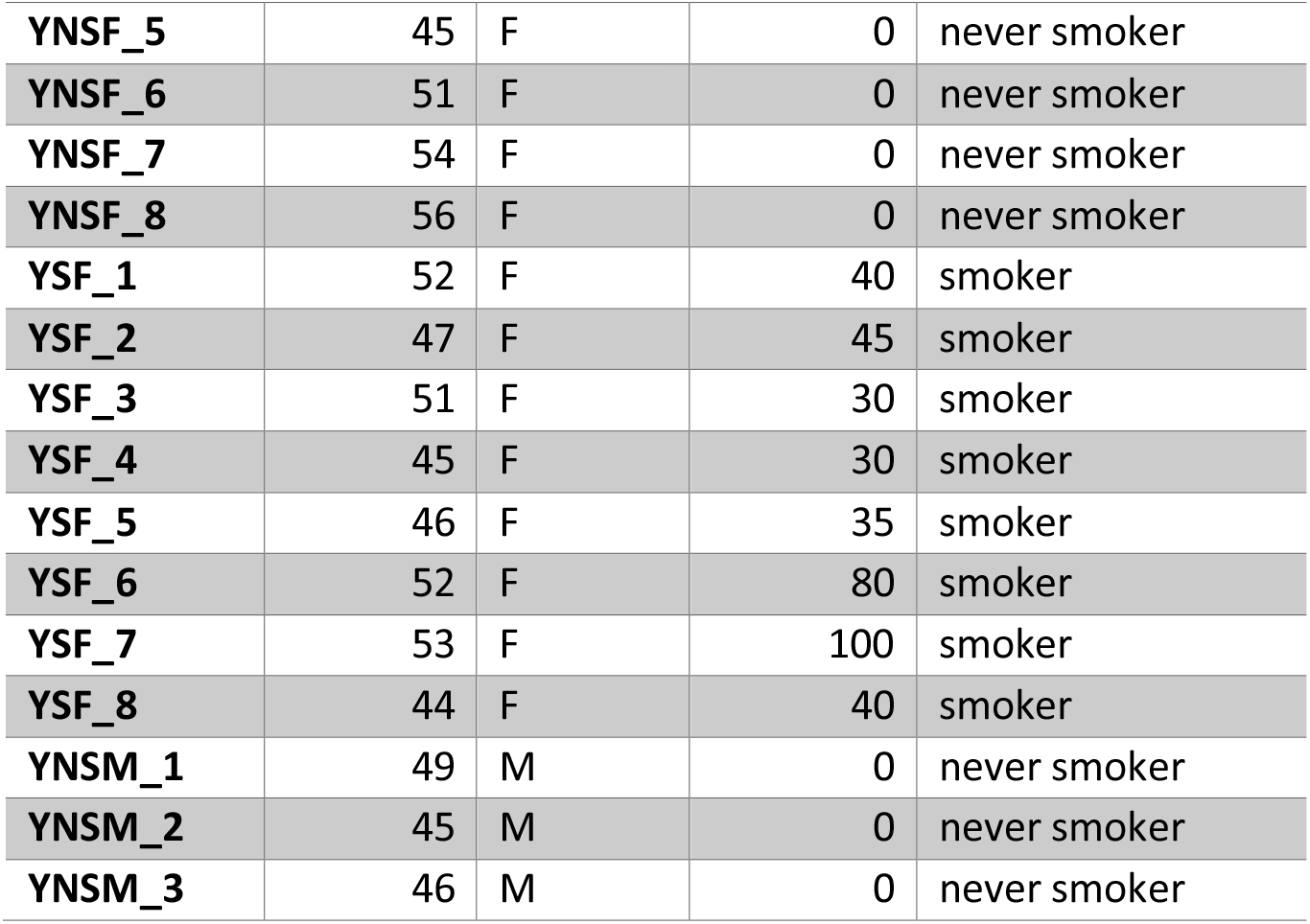
Patient information.

| **pseudonym** | **age** | **sex** | **packyears** | **Smoking habit** |
| --- | --- | --- | --- | --- |
| ENSF_1 | 75 | F | 0 | never smoker |
| ENSF_2 | 78 | F | 0 | never smoker |
| ENSF_3 | 79 | F | 0 | never smoker |
| ENSF_4 | 85 | F | 0 | never smoker |
| ENSF_5 | 88 | F | 0 | never smoker |
| ENSF_6 | 76 | F | 0 | never smoker |
| ENSF_7 | 79 | F | 0 | never smoker |
| YNSF_1 | 50 | F | 0 | never smoker |
| YNSF_2 | 55 | F | 0 | never smoker |
| YNSF_3 | 40 | F | 0 | never smoker |
| YNSF_4 | 57 | F | 0 | never smoker |
| YNSF_5 | 45 | F | 0 | never smoker |
| YNSF_6 | 51 | F | 0 | never smoker |
| YNSF_7 | 54 | F | 0 | never smoker |
| YNSF_8 | 56 | F | 0 | never smoker |
| YSF_1 | 52 | F | 40 | smoker |
| YSF_2 | 47 | F | 45 | smoker |
| YSF_3 | 51 | F | 30 | smoker |
| YSF_4 | 45 | F | 30 | smoker |
| YSF_5 | 46 | F | 35 | smoker |
| YSF_6 | 52 | F | 80 | smoker |
| YSF_7 | 53 | F | 100 | smoker |
| YSF_8 | 44 | F | 40 | smoker |
| YNSM_1 | 49 | M | 0 | never smoker |
| YNSM_2 | 45 | M | 0 | never smoker |
| YNSM_3 | 46 | M | 0 | never smoker |

**Supplementary Table 3 – DE healthy tissue cell type.** Differentially expressed genes with a p-value < 0.01 for all identified clusters in healthy tissue.

**Supplementary Table 4 – DE tumour micoenvironment cell identity.** Differentially expressed genes with a p-value < 0.01 for all identified clusters in tumour originating tissue.

**Supplementary Table 5 – DE malignant cell identity.** Differentially expressed genes with a p-value < 0.01 for cells identified as malignant.

**Supplementary Table 6 –.** GO signatures. GO annotations as in Fig. 3C with corresponding signature annotation

| IDs | GO TERMS | Signature |
| --- | --- | --- |
| 1 | GO_ORGANELLE_FISSION | Mitotic |
| 2 | GO_MITOTIC_NUCLEAR_DIVISION | Mitotic |
| 3 | GO_CHROMOSOMAL_REGION | Mitotic |
| 4 | GO_CONDENSED_CHROMOSOME | Mitotic |
| 5 | GO_CHROMOSOME_CENTROMERIC_REGION | Mitotic |
| 6 | GO_CELL_CYCLE_G2_M_PHASE_TRANSITION | Mitotic |
| 7 | GO_REGULATION_OF_CELL_CYCLE_G2_M_PHASE_TRANSITION | Mitotic |
| 8 | GO_SMALL_GTPASE_BINDING | Mitotic |
| 9 | GO_ANAPHASE_PROMOTING_COMPLEX_DEPENDENT_CATABOLIC_PROCESS | Respiration |
| 10 | GO_ATP_SYNTHESIS_COUPLED_ELECTRON_TRANSPORT | Respiration |
| 11 | GO_RESPIRATORY_ELECTRON_TRANSPORT_CHAIN | Respiration |
| 12 | GO_ATP_METABOLIC_PROCESS | Respiration |
| 13 | GO_OXIDATIVE_PHOSPHORYLATION | Respiration |
| 14 | GO_RESPIRASOME | Respiration |
| 15 | GO_RESPIRATORY_CHAIN_COMPLEX | Respiration |
| 16 | GO_PROTEIN_LOCALIZATION_TO_ENDOPLASMIC_RETICULUM | Transcription |
| 17 | GO_ESTABLISHMENT_OF_PROTEIN_LOCALIZATION_TO_ENDOPLASMIC_RETICULUM | Transcription |
| 18 | GO_COTRANSLATIONAL_PROTEIN_TARGETING_TO_MEMBRANE | Transcription |
| 19 | GO_CYTOSOLIC_RIBOSOME | Transcription |
| 20 | GO_NUCLEAR_TRANSCRIBED_MRNA_CATABOLIC_PROCESS_NONSENSE_MEDIATED_DECAY | Transcription |
| 21 | GO_CELL_ADHESION_MEDIATED_BY_INTEGRIN | Adhesion |
| 22 | GO_RESPONSE_TO_MOLECULE_OF_BACTERIAL_ORIGIN | Adhesion |
| 23 | GO_PROTEIN_COMPLEX_INVOLVED_IN_CELL_ADHESION | Adhesion |
| 24 | GO_INTEGRIN_BINDING | Adhesion |
| 25 | GO_EXOGENOUS_PROTEIN_BINDING | Adhesion |
| 26 | GO_ORGANIC_ACID_TRANSPORT | Metabolism |
| 27 | GO_UNSATURATED_FATTY_ACID_METABOLIC_PROCESS | Metabolism |
| 28 | GO_ICOSANOID_METABOLIC_PROCESS | Metabolism |
| 29 | GO_PROSTANOID_METABOLIC_PROCESS | Metabolism |
| 30 | GO_MONOCARBOXYLIC_ACID_TRANSPORT | Metabolism |
| 31 | GO_NEURON_PROJECTION_ARBORIZATION | Morphology |
| 32 | GO_ACTIN_BASED_CELL_PROJECTION | Morphology |
| 33 | GO_CELL_LEADING_EDGE | Morphology |
| 34 | GO_REGULATION_OF_CELL_MORPHOGENESIS | Morphology |
| 35 | GO_SYNAPSE_ORGANIZATION | Morphology |
| 36 | GO_ACTIN_BINDING | Morphology |
| 37 | GO_PHOSPHOLIPID_BINDING | Phospholipid Binding |
| 38 | GO_RESPONSE_TO_IMMOBILIZATION_STRESS | Phospholipid Binding |
| 39 | GO_PHOSPHATIDYLINOSITOL_BINDING | Phospholipid Binding |
| 40 | GO_MHC_CLASS_II_PROTEIN_COMPLEX | Immune Related |
| 41 | GO_LEUKOCYTE_PROLIFERATION | Immune Related |
| 42 | GO_REGULATION_OF_LEUKOCYTE_PROLIFERATION | Immune Related |
| 43 | GO_T_CELL_PROLIFERATION | Immune Related |
| 44 | GO_PHAGOCYTOSIS | Immune Related |

**Supplementary Table 7 – DE trajectory branches.** Differentially expressed genes with a p-value < 0.01 for cells identified by trajectory branch.

**Supplementary Table 8 – top factor defining genes.** Gene usage per factor determined by NMF.

## Acknowledgments

The authors would like to thank the tissue donors and their families;

David Ibberson (Heidelberg University), Ulrike Krüger (Berlin Institute of Health [BIH], Berlin), and Marten Jager (BIH, Berlin) for next-generation sequencing services, Christa Stolp and Martin Fallenbüchel (Thoraxklinik-Heidelberg) for tissue collection, Luca Tosti (BIH, Berlin) for helpful discussions and Teresa Krieger (BIH, Berlin) for critically revising the manuscript.

This study was supported by the Human Cell Atlas studies of the Chan Zuckerberg initiative and the German Center for Lung Research (DZL, grant number 82DZL00402).

## Author contributions

T.T., M.A.S. and C.C. conceived and designed the study; M.M., C.C., T.M. and R.E. supervised the study; T.T. performed scRNA-Seq experiments; M.A.S. performed histochemical staining; K.J. and L.C. assisted with experiments; T.T. analysed the data; H.W. and M.K. provided tissues and pathological assessments. All authors revised and approved the manuscript.

## Material and Methods

### Sample procurement

Cryopreserved surgical lung tissue from patients with lung adenocarcinoma was provided by the Lung Biobank Heidelberg. All subjects gave their informed consent for inclusion before participation in the study.

This study was conducted in accordance with the Declaration of Helsinki and the Department of Health and Human Services Belmont Report. The use of biomaterial for this study was approved by the local ethics committee of the Medical Faculty Heidelberg (S-270/2001 (biobank vote) and S-056/2021 (study vote)).

Tumour tissue and an additional representative part of normal lung tissue distant from the tumour (> 5 cm) was collected during routine surgical intervention. Pieces of 0.5-1 cm^3^ were cut immediately after resection snap-frozen in liquid nitrogen within 30 min after resection, with no direct contact of samples and nitrogen. After snap-frezing, the vials were stored at - 80°C and monitored regarding temperature until use.

### Single nucleus RNA sequencing

Single nuclei were prepared from frozen tissue as described in [81]. Briefly, snap-frozen healthy lung tissue from lung adenocarcinoma patients was cut into pieces with less than 0.3 cm diameter and single nuclei were isolated at low pH by homogenizing the cells in 1 ml of citric acid-based buffer (Sucrose 0.25 M, Citric Acid 25 mM, Hoechst 33342 1 g/ml) at 4°C using a glass Dounce tissue grinder. After one stroke with the “loose” pestle, the tissue was incubated for 5 minutes on ice, further homogenized with 3–5 strokes of the “loose” pestle, followed by another incubation at 4°C for 5 min. Nuclei were released by five additional strokes with the “tight” pestle, and the nuclei solution was filtered through a 35-µm cell strainer. Cell debris was removed via centrifugation at 4°C for 5 min at 500 g, and the supernatant was removed, followed by nuclei cell pellet resuspension in 700 µl citric acid-based buffer and centrifugation at 4°C for 5 min at 500 g. After carefully removing the supernatant, the nuclei cell pellet was resuspended in 100 µl cold resuspension buffer (25 mM KCl, 3 mM MgCl2, 50 mM Tris-buffer, 400 U RNaseIn, 1 mM DTT, 400 U SUPERase In (AM2694, Thermo Fisher Scientific), 1 g/ml Hoechst (H33342, Thermo Fisher Scientific)). Nuclei concentration was determined using the Countess II FL Automated Cell Counter, and optimal nuclei concentration was obtained by adding additional cold resuspension buffer, if needed. Subsequently, samples were processed using the 10× Chromium device with the 10× Genomics scRNA-Seq protocol v2 to generate cell and gel bead emulsions, followed by reverse transcription, cDNA amplification, and sequencing library preparation following the manufacturers’ instructions. Libraries have subsequently been sequenced one sample per lane on HiSeq4000 (Illumina; paired-end 26 × 74 bp)

### Data pre-processing

For alignment of raw sequencing reads the CellRanger software version 2.1.1 (10x Genomics) together with the human reference genome hg19 was used. Low-quality cells were removed using Seurat version 3.1.3 [27] based on the following on detected genes, RNA molecules and mitochondrial reads (discarded were cells with: genes detected < 200 or, depending on the sample > 3000-10000; molecule count > 7000-90000; and < 10% mitochondrial reads).

Remaining cells were further processed using the Seurat software for log-normalisation, scaling, clustering and UMAP visualisation. Afterwards all healthy control samples were merged using the “FindIntegrationAnchors” and “IntegrateData” functions. Tumour derived samples were then merged with the integrated control data set by the same method, using control data as reference.

### Cell type assignment

Cell types in healthy samples were assigned using canonical marker expression. Subsequently cells of the tumour microenvironment were identified using the “FindTransferAnchors” and “TransferData” functions of the Seurat software, using healthy control data as reference. Similarity scores with reference cell types combined with canonical marker expression were used for cell type assignment. Tumour cell cluster with low similarity scores to reference data and ambiguous marker expression were marked as unidentified.

### Differential expression and gene set enrichment

Cluster specific gene expression was calculated using Wilcoxon-rank-sum-test as implemented in the “FindMarker” function of Seurat. Only genes with adjusted p-value below 0.01 were considered for further analysis. Gene set enrichment was performed using clusterProfiler [82] and version 7.2 of Gene Ontology terms [83].

To decrease the influence of interpatient heterogeneity, when comparing in a given cluster cells from smokers and never smokers, differential gene expression analysis was performed for each patient separate against all other cells within the same cluster from patients of the other smoking group. Only genes that where differentially expressed in at least 40% of patients in each respective group were considered for further analysis.

### CNV inference

To infer copy number variations from single cell transcriptome data the method implemented in the inferCNV software was employed [84].Briefly, patient raw counts of matched healthy control samples were used as baseline expression for an aggregate of genes in proximal genomic location and compared to average expression levels of genes from tumour samples at the same genomic location. Higher average expression in tumour samples was used as indication for copy number gain, while lower expression was indicative of copy number loss at a given genomic location.

### Cell cell communication

For interrogating cell to cell communication the curated database of ligand-receptor pairs and statistical framework as implemented in CellPhoneDB [30] was used. Raw counts for each cell were normalised by library size and mean expression for each gene in the database is calculated. Cells are pooled by cell cluster annotation and the percentage of cells in this cluster expressing each gene is assesed. Through random shuffling of cell labels a null distribution for each gene pair is derived, taking into account the expression levels. This is compared to the observed mean expression of ligand and receptor in two clusters of cells and a p-value for their expression, specifically in this pair of clusters derived from the null distribution. Ligand-receptor pairs are then ranked by p-value and significant interactions are determined.

### Transcription factor networks

Transcription factor networks were inferred using a three-step algorithm implemented in the SCENIC software [34]. First a set of co-expressed genes for each transcription factor is established using GRNBost2. Genes in each set are then filtered by positive correlation with the transcription factor binding motif and scored for their importance in each cell, by the AUCell method further described in the SCENIC software documentation.

### Trajectory analysis

Inference of a possible developmental trajectory was realised using a framework employing principal graph inference, published as STREAM [42]. Briefly, the integrated and normalised expression matrix of all malignant cells is used to first define variable genes using non-parametric local regression, which are the used to reduce the dimensionality of the data employing modified locally linear embedding. This method provides a continuous embedding, by considering local similarity to its neighbours. In this space cells are cluster employing the affinity propagation method. The result is then used to construct a minimum spanning tree to use an initial tree structure for the construction of an elastic principal graph.

### Non negative matrix factorisation

Matrix factorisation was conducted with the use of algorithms implemented in the NNLM software [85]. The matrix of integrated, normalised gene counts and cells was decomposed in two matrices with one fixed dimension, the factor number. Contribution of either a factor to a cell or a gene to a factor was calculated on the appropriate decomposed matrix.

### Immunohistochemical stainings

Paraffin-embedded tissue sections were deparaffinized and peroxidases were blocked for 10 min at room temperature (RT) using 3 % H_2_O_2_ (Applichem, Darmstadt, Germany). Antigen retrieval was performed in a steamer with sodium-citrate-buffer (10 mM sodium citrate, 0.05% Tween 20, pH 6.0) for 15 min. The staining procedure for the polyclonal anti-glycodelin antibody (sc-12289, Santa Cruz Biotechnology, Heidelberg, Germany) was performed with DAKO EnVision+ System-HRP (AEC) for rabbit primary antibodies (Dako, Hamburg, Germany). The tissue slides were incubated overnight at 4°C with an anti-glycodelin antibody at a concentration of 2.5 µg/ml. A linker (rabbit anti-goat IgG, A27001, Thermo Scientific) antibody was used for 30 min at room temperature before tissue sections were incubated with secondary antibody for another 30 min at room temperature. Visualization of glycodelin was performed with AEC+ Substrate-Chromogen (Dako). For ANXA1 staining, the staining procedure was performed with SignalStain® DAB Substrate Kit (#8059, Cell Signaling, Frankfurt, Germany) according to manufacturer’s instructions, using rabbit polyclonal anti-ANXA1 antibody (#32934, Cell Signaling, Frankfurt, Germany). Cell nuclei were stained using Mayer’s Hematoxylin Solution (Sigma-Aldrich, Munich, Germany). Staining was observed with an Olympus IX-71 inverted microscope. Pictures were taken with an Olympus Color View II digital camera and Olympus Cell-F software (cellSense dimension, V1.11, Olympus, Hamburg, Germany). Tiffs were assembled into figures using Photoshop CS6 (Adobe, San José, CA, USA). Only changes in brightness and contrast were applied. Scoring was performed by multiplication of staining intensity (0-3) with the proportion of positive cells (0-4). For each patient, five randomly selected pictures were analysed and median was calculated.

### Statistical analysis and visualisation

Statistical analysis and visualisation has been performed using python 3.7 or R 3.6.3 together with beforementioned software, ggplot2 and ComplexHeatmaps [86].

## Data availability

Raw sequencing access-protected data on the European Genome-Phenome Archive are available at https://www.ebi.ac.uk/ega/home under EGAXXX. Count and sample meta data are available on https://figshare.com/XXX and code describing the analysis on https://github.com/XXX.

## References

1. Islami, F., et al., Proportion and number of cancer cases and deaths attributable to potentially modifiable risk factors in the United States. CA Cancer J Clin, 2018. 68(1): p. 31–54.

2. Sung, H., et al., Global cancer statistics 2020: GLOBOCAN estimates of incidence and mortality worldwide for 36 cancers in 185 countries. CA Cancer J Clin, 2021.

3. Torre, L.A., R.L. Siegel, and A. Jemal, Lung Cancer Statistics. Adv Exp Med Biol, 2016. 893: p. 1–19.

4. Youlden, D.R., S.M. Cramb, and P.D. Baade, The International Epidemiology of Lung Cancer: geographical distribution and secular trends. J Thorac Oncol, 2008. 3(8): p. 819–31.

5. Parkin, D.M., et al., Global cancer statistics, 2002. CA Cancer J Clin, 2005. 55(2): p. 74–108.

6. Wakelee, H.A., et al., Lung cancer incidence in never smokers. J Clin Oncol, 2007. 25(5): p. 472–8.

7. Alberg, A.J., et al., Epidemiology of lung cancer: Diagnosis and management of lung cancer, 3rd ed: American College of Chest Physicians evidence-based clinical practice guidelines. Chest, 2013. 143(5 Suppl): p. e1S–e29S.

8. Bilano, V., et al., Global trends and projections for tobacco use, 1990-2025: an analysis of smoking indicators from the WHO Comprehensive Information Systems for Tobacco Control. Lancet, 2015. 385(9972): p. 966–76.

9. Organization, W.H., WHO report on the global tobacco epidemic 2019: offer help to quit tobacco use. 2019: World Health Organization.

10. Forey, B. and P.N. Lee, New edition of International Smoking Statistics. Int J Epidemiol, 2007. 36(2): p. 471–2.

11. Zheng, M., Classification and Pathology of Lung Cancer. Surg Oncol Clin N Am, 2016. 25(3): p. 447–68.

12. Travis, W.D., et al., Introduction to The 2015 World Health Organization Classification of Tumors of the Lung, Pleura, Thymus, and Heart. J Thorac Oncol, 2015. 10(9): p. 1240–1242.

13. Cancer Genome Atlas Research, N., Comprehensive molecular profiling of lung adenocarcinoma. Nature, 2014. 511(7511): p. 543–50.

14. Kim, N., et al., Single-cell RNA sequencing demonstrates the molecular and cellular reprogramming of metastatic lung adenocarcinoma. Nat Commun, 2020. 11(1): p. 2285.

15. Maynard, A., et al., Therapy-Induced Evolution of Human Lung Cancer Revealed by Single-Cell RNA Sequencing. Cell, 2020. 182(5): p. 1232–1251 e22.

16. Ma, K.Y., et al., Single-cell RNA sequencing of lung adenocarcinoma reveals heterogeneity of immune response-related genes. JCI Insight, 2019. 4(4).

17. Lavin, Y., et al., Innate Immune Landscape in Early Lung Adenocarcinoma by Paired Single-Cell Analyses. Cell, 2017. 169(4): p. 750–765 e17.

18. Chen, J., et al., Single-cell transcriptome and antigen-immunoglobin analysis reveals the diversity of B cells in non-small cell lung cancer. Genome Biol, 2020. 21(1): p. 152.

19. Sinjab, A., et al., Resolving the spatial and cellular architecture of lung adenocarcinoma by multi-region single-cell sequencing. bioRxiv, 2020: p. 2020.09.04.283739.

20. Lambrechts, D., et al., Phenotype molding of stromal cells in the lung tumor microenvironment. Nat Med, 2018. 24(8): p. 1277–1289.

21. Li, B., et al., Comprehensive analyses of tumor immunity: implications for cancer immunotherapy. Genome Biol, 2016. 17(1): p. 174.

22. Lim, S.M., M.H. Hong, and H.R. Kim, Immunotherapy for Non-small Cell Lung Cancer: Current Landscape and Future Perspectives. Immune Netw, 2020. 20(1): p. e10.

23. Waldman, A.D., J.M. Fritz, and M.J. Lenardo, A guide to cancer immunotherapy: from T cell basic science to clinical practice. Nat Rev Immunol, 2020. 20(11): p. 651–668.

24. Singh, N., et al., Inflammation and cancer. Ann Afr Med, 2019. 18(3): p. 121–126.

25. Wang, D. and R.N. DuBois, Immunosuppression associated with chronic inflammation in the tumor microenvironment. Carcinogenesis, 2015. 36(10): p. 1085–93.

26. Lukassen, S., et al., SARS-CoV-2 receptor ACE2 and TMPRSS2 are primarily expressed in bronchial transient secretory cells. EMBO J, 2020. 39(10): p. e105114.

27. Stuart, T., et al., Comprehensive Integration of Single-Cell Data. Cell, 2019. 177(7): p. 1888–1902 e21.

28. Todisco, T., et al., Normal reference values for regional pulmonary peripheral airspace epithelial permeability. Influence of pneumonectomy and the smoking habit. Respiration, 1989. 55(2): p. 84–93.

29. Wright, J.L., et al., Airway inflammation and peribronchiolar attachments in the lungs of nonsmokers, current and ex-smokers. Lung, 1988. 166(5): p. 277–86.

30. Efremova, M., et al., CellPhoneDB: inferring cell-cell communication from combined expression of multi-subunit ligand-receptor complexes. Nat Protoc, 2020. 15(4): p. 1484–1506.

31. Negrini, S., V.G. Gorgoulis, and T.D. Halazonetis, Genomic instability--an evolving hallmark of cancer. Nat Rev Mol Cell Biol, 2010. 11(3): p. 220–8.

32. Stranger, B.E., et al., Relative impact of nucleotide and copy number variation on gene expression phenotypes. Science, 2007. 315(5813): p. 848–53.

33. Lawson, D.A., et al., Tumour heterogeneity and metastasis at single-cell resolution. Nat Cell Biol, 2018. 20(12): p. 1349–1360.

34. Aibar, S., et al., SCENIC: single-cell regulatory network inference and clustering. Nat Methods, 2017. 14(11): p. 1083–1086.

35. Singleton, D.C. and A.L. Harris, Targeting the ATF4 pathway in cancer therapy. Expert Opin Ther Targets, 2012. 16(12): p. 1189–202.

36. Harding, H.P., et al., An integrated stress response regulates amino acid metabolism and resistance to oxidative stress. Mol Cell, 2003. 11(3): p. 619–33.

37. Takeda, J., S. Seino, and G.I. Bell, Human Oct3 gene family: cDNA sequences, alternative splicing, gene organization, chromosomal location, and expression at low levels in adult tissues. Nucleic Acids Res, 1992. 20(17): p. 4613–20.

38. Niwa, H., J. Miyazaki, and A.G. Smith, Quantitative expression of Oct-3/4 defines differentiation, dedifferentiation or self-renewal of ES cells. Nat Genet, 2000. 24(4): p. 372–6.

39. Yu, W., et al., MEF2 transcription factors promotes EMT and invasiveness of hepatocellular carcinoma through TGF-beta1 autoregulation circuitry. Tumour Biol, 2014. 35(11): p. 10943–51.

40. Busygina, V., et al., Multiple endocrine neoplasia type 1 interacts with forkhead transcription factor CHES1 in DNA damage response. Cancer Res, 2006. 66(17): p. 8397–403.

41. Burrell, R.A., et al., The causes and consequences of genetic heterogeneity in cancer evolution. Nature, 2013. 501(7467): p. 338–45.

42. Chen, H., et al., Single-cell trajectories reconstruction, exploration and mapping of omics data with STREAM. Nat Commun, 2019. 10(1): p. 1903.

43. Lee, J., S. Giordano, and J. Zhang, Autophagy, mitochondria and oxidative stress: cross-talk and redox signalling. Biochem J, 2012. 441(2): p. 523–40.

44. Maman, S. and I.P. Witz, A history of exploring cancer in context. Nat Rev Cancer, 2018. 18(6): p. 359–376.

45. Cheng, G., et al., Identification of PLXDC1 and PLXDC2 as the transmembrane receptors for the multifunctional factor PEDF. Elife, 2014. 3: p. e05401.

46. Martinez-Marin, D., et al., PEDF increases the tumoricidal activity of macrophages towards prostate cancer cells in vitro. PLoS One, 2017. 12(4): p. e0174968.

47. Atkinson, P.G. and C.H. Barton, Ectopic expression of Nramp1 in COS-1 cells modulates iron accumulation. FEBS Lett, 1998. 425(2): p. 239–42.

48. La Fleur, L., et al., Expression of scavenger receptor MARCO defines a targetable tumor-associated macrophage subset in non-small cell lung cancer. Int J Cancer, 2018. 143(7): p. 1741–1752.

49. Georgoudaki, A.M., et al., Reprogramming Tumor-Associated Macrophages by Antibody Targeting Inhibits Cancer Progression and Metastasis. Cell Rep, 2016. 15(9): p. 2000–11.

50. Jiang, Z., et al., Targeting the SLIT/ROBO pathway in tumor progression: molecular mechanisms and therapeutic perspectives. Ther Adv Med Oncol, 2019. 11: p. 1758835919855238.

51. Zhang, C., et al., Effects of Slit3 silencing on the invasive ability of lung carcinoma A549 cells. Oncol Rep, 2015. 34(2): p. 952–60.

52. Nissen, N.I., M. Karsdal, and N. Willumsen, Collagens and Cancer associated fibroblasts in the reactive stroma and its relation to Cancer biology. J Exp Clin Cancer Res, 2019. 38(1): p. 115.

53. Zou, X., et al., Up-regulation of type I collagen during tumorigenesis of colorectal cancer revealed by quantitative proteomic analysis. J Proteomics, 2013. 94: p. 473–85.

54. Willumsen, N., et al., Serum biomarkers reflecting specific tumor tissue remodeling processes are valuable diagnostic tools for lung cancer. Cancer Med, 2014. 3(5): p. 1136–45.

55. Hirai, K., et al., The spread of human lung cancer cells on collagens and its inhibition by type III collagen. Clin Exp Metastasis, 1991. 9(6): p. 517–27.

56. Bedard, P.L., et al., Tumour heterogeneity in the clinic. Nature, 2013. 501(7467): p. 355–64.

57. Dvorak, H.F., Tumors: wounds that do not heal-redux. Cancer Immunol Res, 2015. 3(1): p. 1–11.

58. Schafer, M. and S. Werner, Cancer as an overhealing wound: an old hypothesis revisited. Nat Rev Mol Cell Biol, 2008. 9(8): p. 628–38.

59. Dvorak, H.F., Tumors: wounds that do not heal. Similarities between tumor stroma generation and wound healing. N Engl J Med, 1986. 315(26): p. 1650–9.

60. Virchow, R., Aetiologie der neoplastischen Geschwulste/Pathogenie der neoplastischen Geschwulste. 1863.

61. Rybinski, B., J. Franco-Barraza, and E. Cukierman, The wound healing, chronic fibrosis, and cancer progression triad. Physiol Genomics, 2014. 46(7): p. 223–44.

62. Rosenfeldt, M.T. and K.M. Ryan, The multiple roles of autophagy in cancer. Carcinogenesis, 2011. 32(7): p. 955–63.

63. Yun, C.W. and S.H. Lee, The Roles of Autophagy in Cancer. Int J Mol Sci, 2018. 19(11).

64. Cirino, G., et al., Recombinant human lipocortin 1 inhibits thromboxane release from guinea-pig isolated perfused lung. Nature, 1987. 328(6127): p. 270–2.

65. Lai, T., et al., Annexin A1 is elevated in patients with COPD and affects lung fibroblast function. Int J Chron Obstruct Pulmon Dis, 2018. 13: p. 473–486.

66. Yang, G., et al., Identification of genes and analysis of prognostic values in nonsmoking females with non-small cell lung carcinoma by bioinformatics analyses. Cancer Manag Res, 2018. 10: p. 4287–4295.

67. Rachmilewitz, J., G.J. Riely, and M.L. Tykocinski, Placental protein 14 functions as a direct T-cell inhibitor. Cell Immunol, 1999. 191(1): p. 26–33.

68. Soni, C. and A.A. Karande, Glycodelin A suppresses the cytolytic activity of CD8+ T lymphocytes. Mol Immunol, 2010. 47(15): p. 2458–66.

69. Ren, S., et al., Functional characterization of the progestagen-associated endometrial protein gene in human melanoma. J Cell Mol Med, 2010. 14(6B): p. 1432–42.

70. Schneider, M.A., et al., Glycodelin: A New Biomarker with Immunomodulatory Functions in Non-Small Cell Lung Cancer. Clin Cancer Res, 2015. 21(15): p. 3529–40.

71. Wanner, A., A review of the effects of cigarette smoke on airway mucosal function. Eur J Respir Dis Suppl, 1985. 139: p. 49–53.

72. Lee, J., V. Taneja, and R. Vassallo, Cigarette smoking and inflammation: cellular and molecular mechanisms. J Dent Res, 2012. 91(2): p. 142–9.

73. Beane, J., et al., Characterizing the impact of smoking and lung cancer on the airway transcriptome using RNA-Seq. Cancer Prev Res (Phila), 2011. 4(6): p. 803–17.

74. Martos, S.N., et al., Single-cell analyses identify tobacco smoke exposure-associated, dysfunctional CD16<sup>+</sup> CD8 T cells with high cytolytic potential in peripheral blood. bioRxiv, 2019: p. 783126.

75. Duclos, G.E., et al., Characterizing smoking-induced transcriptional heterogeneity in the human bronchial epithelium at single-cell resolution. Sci Adv, 2019. 5(12): p. eaaw3413.

76. Goldfarbmuren, K.C., et al., Dissecting the cellular specificity of smoking effects and reconstructing lineages in the human airway epithelium. Nat Commun, 2020. 11(1): p. 2485.

77. Brasky, T.M., et al., Non-steroidal anti-inflammatory drugs and small cell lung cancer risk in the VITAL study. Lung Cancer, 2012. 77(2): p. 260–4.

78. Zappavigna, S., et al., Anti-Inflammatory Drugs as Anticancer Agents. Int J Mol Sci, 2020. 21(7).

79. Young, R.P., et al., COPD prevalence is increased in lung cancer, independent of age, sex and smoking history. Eur Respir J, 2009. 34(2): p. 380–6.

80. Durham, A.L. and I.M. Adcock, The relationship between COPD and lung cancer. Lung Cancer, 2015. 90(2): p. 121–7.

81. Tosti, L., et al., Single-Nucleus and In Situ RNA-Sequencing Reveal Cell Topographies in the Human Pancreas. Gastroenterology, 2021. 160(4): p. 1330–1344 e11.

82. Yu, G., et al., clusterProfiler: an R package for comparing biological themes among gene clusters. OMICS, 2012. 16(5): p. 284–7.

83. Gene Ontology, C., The Gene Ontology resource: enriching a GOld mine. Nucleic Acids Res, 2021. 49(D1): p. D325–D334.

84. Tickle, T., et al., inferCNV of the Trinity CTAT Project. 2019.

85. Lin, X. and P.C. Boutros, Optimization and expansion of non-negative matrix factorization. BMC Bioinformatics, 2020. 21(1): p. 7.

86. Gu, Z., R. Eils, and M. Schlesner, Complex heatmaps reveal patterns and correlations in multidimensional genomic data. Bioinformatics, 2016. 32(18): p. 2847–9.

